# Transgenic mice overexpressing mutant TDP-43 show aberrant splicing of autism associated gene *Zmynd11* prior to onset of motor symptoms

**DOI:** 10.1101/2022.12.11.519991

**Authors:** Ramesh K. Narayanan, Ajay Panwar, Tim J. Butler, Anthony N. Cutrupi, Marina Kennerson, Steve Vucic, Ashokkumar Balasubramaniem, Marie Mangelsdorf, Robyn H. Wallace

## Abstract

Mutations in TDP-43 are known to cause Amyotrophic Lateral Sclerosis (ALS) and Frontotemporal Dementia (FTD). TDP-43 binds to and regulates splicing of several RNA including *Zmynd11*. Zmynd11 is a transcriptional repressor and a potential E3 ubiquitin ligase family member, known for its role in neuron and muscle differentiation. Mutations in *Zmynd11* have been associated with autism with significant developmental motor delays, intellectual disability, and ataxia. Here, we show that *Zmynd11* is aberrantly spliced in the brain and spinal cord of transgenic mice overexpressing a mutant human TDP-43 (A315T), and that these changes occur before the onset of motor symptoms.

## Description

The discovery of TDP-43 as part of cytoplasmic inclusions in ALS patient spinal cord motor neurons (Neumann, Sampathu et al. 2006) and the subsequent discovery of mutations in the *TARDBP* gene that encodes TDP-43 (Sreedharan, Blair et al. 2008) revolutionized the field of ALS research. TDP-43 is a nucleo-cytoplasmic shuttling protein that binds to target mRNA and regulates their transport and/or splicing (Sephton, Cenik et al. 2010, Tollervey, Curk et al. 2011, Colombrita, Onesto et al. 2012, Narayanan 2013, Narayanan, Mangelsdorf et al. 2013). In the first mouse model of TDP-43 developed, Prp-TDP43^A315T^, animals were engineered to express the human mutant gene under the control of a prion protein promoter driving expression predominantly in the mouse CNS. The transgenic mice developed symptoms around 12 weeks of age and displayed features of ALS and FTD (Wegorzewska, Bell et al. 2009). Therefore, for the purpose of this study, 50-day old (pre-symptomatic) and 100-day old animals (post-symptomatic) were classified into respective groups, to represent stages of the disease. Among the aberrantly spliced genes, zinc finger, MYND type-containing 11 (*Zmynd11*) is of particular interest due to its recent implications in autism-related motor delay (Moskowitz, Belnap et al. 2016); intellectual disability (Pruccoli, Graziano et al. 2021) and brain atrophy and ataxia (Indelicato, Zech et al. 2022). In addition, *Zmynd11/ZMYND11* has been previously reported to be an RNA binding partner of TDP-43 in the mouse brain (Narayanan, Mangelsdorf et al. 2013), rat cortical neurons (Sephton, Cenik et al. 2010) and post-mortem human brain (Tollervey, Curk et al. 2011) and was reported to be aberrantly spliced in the spinal cord of sporadic ALS patients with TDP-43 pathology (Rabin, Kim et al. 2010).

To identify aberrant splicing events, total RNA isolated from the brain and spinal cord of pre- and post-symptomatic transgenic Prp-TDP43^A315T^ (Tg) animals and their age-matched wild type (Wt) controls were hybridized to Affymetrix mouse exon 1.0 ST arrays. Our microarray analysis identified several significant disease-stage specific splicing events in the brain and spinal cord of Tg animals (Array Express Acc. No. E-MTAB-1487). The complete list of aberrantly spliced genes in the mouse CNS from this study can be found here (Narayanan 2013). Twenty-six genes were identified to be to aberrantly spliced, in both brain and spinal cord at pre- and post-symptomatic disease stages. *Zmynd11* was among them and predicted to be aberrantly spliced in the brain of pre-symptomatic (alt-splice p value - 2.4 × 10^−03^) and post-symptomatic (alt-splice p value - 4.3 × 10^−07^) Tg animals. Similarly, *Zmynd11* was predicted to be aberrantly spliced in the spinal cord of pre-symptomatic (p value = 8.5 ×10^−03^) and post-symptomatic (p value = 9.5 × 10^−03^) Tg animals. Microarray analysis showed that two *Zmynd11* transcripts encoding a longer gamma isoform (GenBank accession number NM_001199141; 4132 bp) and a shorter alpha isoform (GenBank accession number NM_144516; 3970 bp), which lacks an exon that codes for a plant homeo domain (PHD) and is aberrantly spliced in Tg animals. The expression trace for individual exons of the *Zmynd11* gene in the brain and spinal cord of Tg animals and their corresponding age matched Wt is shown in Figure 1 (A and B). Decreased expression of exon 4 is indicated by a blue arrow. There was no significant difference in the expression traces for other *Zmynd11* exons between Tg and Wt animals suggesting that the change in exon expression in the predicted region (blue arrow) is due to splicing and not due to a change in overall gene expression.

**Figure 1.**
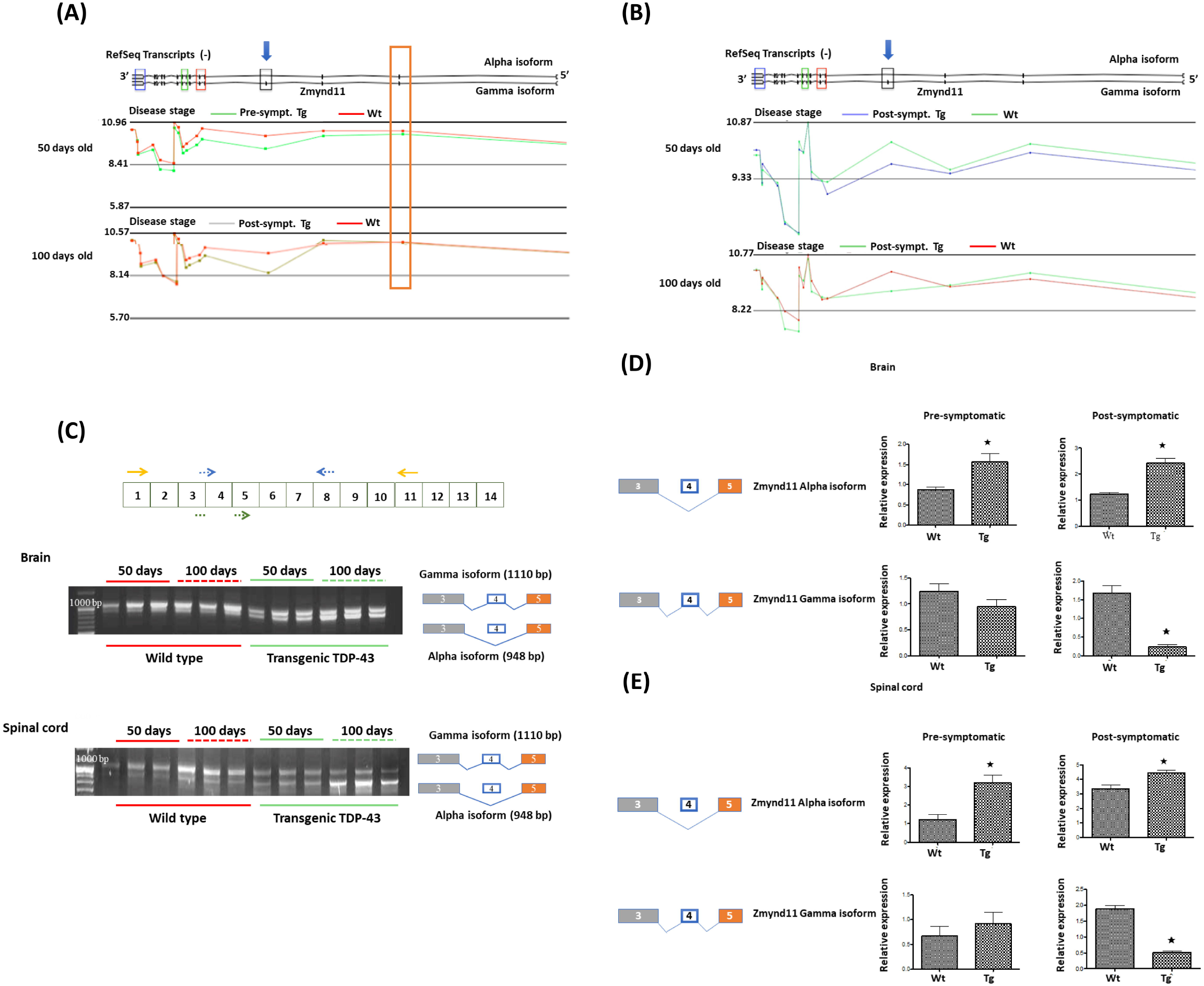
Chromosome view of the microarray data from Partek software showing the expression trace for *Zmynd11* exons in the brain **(A)** and spinal cord **(B)** of pre and post-symptomatic transgenic Prp-TDP43^A315T^ animals and age-matched wild type controls; the expression levels are shown on a log2 scale. The blue arrow indicates the exon predicted to be aberrantly spliced. The elongated orange box **(A)** indicates an exon that is not aberrantly spliced. *Zmynd11* exons that code for functional domains are indicated using small rectangular coloured boxes. Red coloured box indicates the region that codes for **Bromodomain**, blue for **Zinc finger myeloid, Nervy, and DEAF-1 domain**, green for **Pro-Trp-Trp-Pro (PWWP)** and black for **plant homeo domain (PHD)** that is aberrantly spliced in Tg animals. **(C)**. Top panel - *Zmynd11* exons and the position of primers used in validation experiments; Orange arrows indicate primers that amplifies both isoforms (Forward:5’-CATGGAGTTCGTGTGTGGAC-3’ and Reverse:5’-CACGTGCAGTCGGTGAACAT-3’); Dotted blue arrow - primers that amplifies gamma isoform (Forward:5’-AGGAGATGAGATTGACTGGG-3’ and Reverse:5’-AATGTCCGCCTGCTCACTGT-3’); Dotted green arrow indicates the forward primer (Forward: CAGGAGATGAGATTAGCATTAAG) when used in combination with reverse primer in exon 8 (dotted blue arrow) amplifies the alpha isoform. Bottom - RT-PCR validation of the *Zmynd11* splicing event identified by microarray in brain and spinal cord of pre and post-symptomatic Tg animals. RT-PCR amplification of the *Zmynd11* isoforms shows a shift in expression towards the shorter isoform (alpha isoform) in the brain and spinal cord tissue of transgenic animals when compared to Wt animals. A schematic representation of the gamma and alpha isoforms is given at the right of the figure, indicating exon 4 is omitted in the alpha isoform. **(D and E)**. Quantification of gamma and alpha isoforms of *Zmynd11* in the brain **(D)** and spinal cord **(E)** of pre- and post-symptomatic Tg animals and their age matched Wt counterparts using qRT-PCR. n = 3 per group; **⋆** = p value < 0.05. Block diagram on the left represents the alpha and gamma isoforms of *Zmynd11* gene. Bar plot on the right shows the quantification results. Expression of the individual isoforms were normalised to *Pgk1* expression (Forward primer: 5’-TGCACGCTTCAAAAGCGCACG-3’ and reverse primer: 5’-AAGTCCACCCTCATCACGACCC-3’). The X-axis indicates the levels of mRNA of *Zmynd11* isoforms relative to the levels of *Pgk1* mRNA. qRT-PCR results show a significant increase in the alpha isoform in the post-symptomatic transgenic animals.

In the Prp-TDP43^A315T^ Tg animal CNS, the *Zmynd11* exon 4 encoding the PHD domain is under expressed potentially leading to an increase in the shorter alpha isoform and consequently a reduction in the full-length gamma isoform. Reverse Transcriptase-PCR (RT-PCR) amplification of *Zmynd11* using primers that amplify both isoforms followed by agarose gel electrophoresis showed a shift in expression towards the alpha isoform in both brain and spinal cord of the pre- and post-symptomatic Tg animals when compared to age-matched Wt littermates (Figure 1C). The shift in expression towards the shorter alpha isoform is evident in pre-symptomatic animals and this shift increased in the post-symptomatic animals. Quantification experiments using isoform-specific primers and qRT-PCR confirmed a significant increase in expression of the shorter alpha isoform in both brain (Figure 1D) and spinal cord tissue (1E) of pre- and post-symptomatic animals Tg animals. The expression of the longer gamma isoform was significantly reduced in the brain and spinal cord of post-symptomatic Tg animals.

*Zmynd11*, initially known as an adenovirus binding protein (Hateboer, Gennissen et al. 1995), is a key epigenetic regulator (Wen, Li et al. 2014) and plays an essential role in inhibition of muscle and neuron differentiation (Yu, Shao et al. 2009). The PHD finger has been previously implicated in chromatin mediated gene regulation (Aasland, Gibson et al. 1995) in addition to playing a role in protein-protein interactions and gene expression (Shi, Hong et al. 2006, Velasco, Grkovic et al. 2006). Furthermore, PHD fingers exhibit E3 ubiquitin ligase activity, a group of enzymes known to play a pivotal role in poly-ubiquitination of protein thus marking them for degradation by proteasome (Dul and Walworth 2007). A comprehensive study of E3 ligase genes reported that variants in 13% of the known E3 ligase genes (~ 80 genes) cause 70 different types of neurological disorders including neurodegenerative diseases such as Parkinsons disease, Spinal Muscular Atrophy and inherited peripheral neuropathies including Charcot-Marie-Tooth Type 2 disease (George, Hoffiz et al. 2018). Imbalance of *Zmynd11* isoforms may lead to abnormal ubiquitination and impaired protein degradation pathways that leads to disrupted protein homeostasis, a signature of neurodegenerative diseases including ALS (Ruegsegger and Saxena 2016, Lambert-Smith, Saunders et al. 2022). Future studies on isoform specific knock down using RNA interference technology in motor neuron-like cells would elucidate the effect of loss of *Zmynd11* full-length isoform on motor neuron function and its contribution to ALS.

We and others have demonstrated that *Zmynd11/ZMYND11* is an RNA target of TDP-43 in mouse/humans and that it is aberrantly spliced in a mouse model of TDP-43 and in ALS patients with TDP-43 pathology. Given its role in global gene regulation, protein homeostasis and involvement in several neurological diseases, we urge other laboratories to screen for the *Zmynd11/ZMYND11* mis-splicing in their TDP-43 mouse models and human post-mortem CNS samples for isoform imbalance. It will be prudent to screen ALS and other neurological diseases patient cohorts for mutations in ZMYND11.

## Methods

### Animals

The University of Queensland Animal Ethics Committee approved all the experiments. Mice overexpressing a mutant human TDP-43 (hTDP-43) gene were obtained from The Jackson Laboratory (strain name - B6;CB-Tg(Prnp-TARDBP*A315T) 95Balo/J; Stock no:010700). C57BL6/J mice were supplied by The University of Queensland Biological Resources department. Animals were euthanised at P50 and P100 (n=3 per genotype) and tissues were harvested immediately for RNA isolation.

### RNA isolation

Brain and spinal cord tissue from Prp-TDP43^A315T^ (Tg) animals and wild type (Wt) littermates were homogenized using a TissueRuptor (Qiagen). 1 mL of TRIzol per 250 mg of tissue was used for homogenization and RNA was isolated as per the manufacturer’s instruction. DNase treatment was carried out using the DNase free kit (Ambion) followed by quantification using Nanodrop.

### Exon array analysis

Total RNA (200 ng) from transgenic TDP-43 mice and control animals was converted to cDNA and then amplified using the Applause WT-Amp Plus ST kit (NuGEN). Amplified cDNA (5 μg) was then fragmented, and biotin labeled using the Encore Biotin Module kit (NuGEN). Labelled cDNA (50 μL) was hybridized onto Affymetrix GeneChip Exon 1.0 ST arrays in a hybridization oven at 45°C for 20 h at 60rpm. Following hybridization, the arrays were washed and stained using Affymetrix fluidics station 450. The GeneChip® scanner was used for scanning labelled arrays and GeneChip® Command Console® Software (AGCC) was used for generating CEL files.

### Microarray data analysis

CEL files were analyzed using the built-in exon array analysis workflow in Partek software. Robust microarray averaging (RMA) was used to normalize the data and Principal component analysis (PCA) was performed to check for sample outliers. Alt-splice ANOVA was used for identifying novel alternative splicing events. A p value ≤ 0.05 was considered significant. Raw and processed data has been submitted to Array Express (Acc.No: E-MTAB-1487).

### Reverse Transcription-PCR (RT-PCR)

cDNA for PCR and qRT-PCR was synthesized using 1 μg of total RNA from the RNA preparation from the brain and spinal cord of Tg and Wt animals used in exon array analysis. cDNA synthesis was carried out using the Superscript™ III First-Strand Synthesis System (Invitrogen) according to the instruction manual.

Reverse transcribed template (1 μL) was mixed with 200 μM dNTPs, 1X ThermoPol buffer, 1.25 U of Taq DNA polymerase, 100 ng of each primer and MilliQ H_2_O in20 μL PCR reaction volume. The PCR was carried out using the DNA Engine Tetrad 2 (Bio-Rad). Cycling conditions for *Zmynd11* RT-PCR were as follows: denaturation at 94°C for 1 min; 10 cycles of denaturation at 94°C for 30 s, annealing at 60°C for 30 s, extension at 72 °C for 30 s; 25 cycles of denaturation at 94°C for 30 s, annealing at 55°C for 30 s and extension at 72°C for 30 s followed by a final extension at 72°C for 2 min. RT-PCR products (10 μL)were size fractionated using a 2% (w/v) agarose gel in 1X TAE buffer containing 25 X SYBR Safe DNA Gel Stain (Invitrogen). Amplified products were visualized under UV light and analyzed using Gel Doc equipment (Bio-Rad) with Quantity One software (Bio-Rad).

### Quantitative Real Time PCR (qRT-PCR)

Quantitative PCR experiments were performed using a LightCycler® 480 (Roche Diagnostics). For qRT-PCR reactions 3 μL of diluted cDNA (1:6) was mixed with 0.4 μM of each primer (forward and reverse) and 1X SYBR Green I Master mix. Cycling conditions for qRT-PCR included a pre-incubation step at 95°C for 5 minutes followed by 45 cycles of 95°C for 10 s, 60 or 64°C for 10 s and a final extension step at 72°C for 30 s. LightCycler® 480 software (release 1.5.0 SP3) was used to calculate Ct values. *Pgk1* was used for gene normalization for qPCR experiments. Statistical analysis for qRT-PCR data was performed using a two-tailed t test with unequal variance. (n=3 per group per genotype).

## Acknowledgement

This work formed of part of RKN’s PhD dissertation and was not published elsewhere. This research was undertaken at the Peter Goodenough and Wantoks Research Lab, Queensland Brain Institute under the supervision of MM and RHW. RHW and MM were supported by the Ross Maclean Fellowship. RKN was supported by The University of Queensland (UQ) tuition fee waiver award, UQ Research Scholarship and QBI top-up scholarship. The work was supported by funds from the MND Research Institute of Australia and the Peter Goodenough estate. The authors declare no conflicting interests.

## Author Contributions

RKN - Conceptualization, experimentation, analysis, Writing – original draft, Writing - Review and Editing; MM and RHW - Conceptualization, supervision, Resources, Writing – Review and Editing; AP and TJB - Animal husbandry and surgery; TJB - Laboratory technical assistance and qPCR; ANC - Array Express data assistance and analysis, and AB - Writing - Review and Editing; MK and SV - Resources and Writing - Review and Editing.

## References

Aasland, R., T. J. Gibson and A. F. Stewart (1995). “The PHD finger: implications for chromatin-mediated transcriptional regulation.” Trends Biochem Sci 20(2): 56–59.

Colombrita, C., E. Onesto, F. Megiorni, A. Pizzuti, F. E. Baralle, E. Buratti, V. Silani and A. Ratti (2012). “TDP-43 and FUS RNA-binding proteins bind distinct sets of cytoplasmic messenger RNAs and differently regulate their post-transcriptional fate in motoneuron-like cells.” The Journal of biological chemistry.

Dul, B. E. and N. C. Walworth (2007). “The plant homeodomain fingers of fission yeast Msc1 exhibit E3 ubiquitin ligase activity.” The Journal of biological chemistry 282(25): 18397–18406.

George, A. J., Y. C. Hoffiz, A. J. Charles, Y. Zhu and A. M. Mabb (2018). “A Comprehensive Atlas of E3 Ubiquitin Ligase Mutations in Neurological Disorders.” Front Genet 9: 29.

Hateboer, G., A. Gennissen, Y. F. Ramos, R. M. Kerkhoven, V. Sonntag-Buck, H. G. Stunnenberg and R. Bernards (1995). “BS69, a novel adenovirus E1A-associated protein that inhibits E1A transactivation.” The EMBO journal 14(13): 3159–3169.

Indelicato, E., M. Zech, M. Amprosi and S. Boesch (2022). “Untangling neurodevelopmental disorders in the adulthood: a movement disorder is the clue.” Orphanet J Rare Dis 17(1): 55.

Lambert-Smith, I. A., D. N. Saunders and J. J. Yerbury (2022). “Proteostasis impairment and ALS.” Prog Biophys Mol Biol.

Moskowitz, A. M., N. Belnap, A. L. Siniard, S. Szelinger, A. M. Claasen, R. F. Richholt, M. De Both, J. J. Corneveaux, C. Balak, I. S. Piras, M. Russell, A. L. Courtright, S. Rangasamy, K. Ramsey, D. W. Craig, V. Narayanan, M. J. Huentelman and I. Schrauwen (2016). “A de novo missense mutation in ZMYND11 is associated with global developmental delay, seizures, and hypotonia.” Cold Spring Harb Mol Case Stud 2(5): a000851.

Narayanan, R. K. (2013). Molecular analysis of the motor neuron disease-associated protein, TDP-43 PhD, The University of Queensland.

Narayanan, R. K., M. Mangelsdorf, A. Panwar, T. J. Butler, P. G. Noakes and R. H. Wallace (2013). “Identification of RNA bound to the TDP-43 ribonucleoprotein complex in the adult mouse brain.” Amyotroph Lateral Scler Frontotemporal Degener 14(4): 252–260.

Neumann, M., D. M. Sampathu, L. K. Kwong, A. C. Truax, M. C. Micsenyi, T. T. Chou, J. Bruce, T. Schuck, M. Grossman, C. M. Clark, L. F. McCluskey, B. L. Miller, E. Masliah, I. R. Mackenzie, H. Feldman, W. Feiden, H. A. Kretzschmar, J. Q. Trojanowski and V. M. Lee (2006). “Ubiquitinated TDP-43 in frontotemporal lobar degeneration and amyotrophic lateral sclerosis.” Science 314(5796): 130–133.

Pruccoli, J., C. Graziano, C. Locatelli, L. Maltoni, H. A. Sheikh Maye and D. M. Cordelli (2021). “Expanding the Neurological Phenotype of Ring Chromosome 10 Syndrome: A Case Report and Review of the Literature.” Genes (Basel) 12(10).

Rabin, S. J., J. M. Kim, M. Baughn, R. T. Libby, Y. J. Kim, Y. Fan, A. La Spada, B. Stone and J. Ravits (2010). “Sporadic ALS has compartment-specific aberrant exon splicing and altered cell-matrix adhesion biology.” Human molecular genetics 19(2): 313–328.

Ruegsegger, C. and S. Saxena (2016). “Proteostasis impairment in ALS.” Brain Res 1648(Pt B): 571–579.

Sephton, C. F., C. Cenik, A. Kucukural, E. B. Dammer, B. Cenik, Y. Han, C. M. Dewey, F. P. Roth, J. Herz, J. Peng, M. J. Moore and G. Yu (2010). “Identification of neuronal RNA targets of TDP-43-containing ribonucleoprotein complexes.” J Biol Chem 286(2): 1204–1215.

Shi, X., T. Hong, K. L. Walter, M. Ewalt, E. Michishita, T. Hung, D. Carney, P. Pena, F. Lan, M. R. Kaadige, N. Lacoste, C. Cayrou, F. Davrazou, A. Saha, B. R. Cairns, D. E. Ayer, T. G. Kutateladze, Y. Shi, J. Cote, K. F. Chua and O. Gozani (2006). “ING2 PHD domain links histone H3 lysine 4 methylation to active gene repression.” Nature 442(7098): 96–99.

Sreedharan, J., I. P. Blair, V. B. Tripathi, X. Hu, C. Vance, B. Rogelj, S. Ackerley, J. C. Durnall, K. L. Williams, E. Buratti, F. Baralle, J. de Belleroche, J. D. Mitchell, P. N. Leigh, A. Al-Chalabi, C. C. Miller, G. Nicholson and C. E. Shaw (2008). “TDP-43 mutations in familial and sporadic amyotrophic lateral sclerosis.” Science 319(5870): 1668–1672.

Tollervey, J. R., T. Curk, B. Rogelj, M. Briese, M. Cereda, M. Kayikci, J. Konig, T. Hortobagyi, A. L. Nishimura, V. Zupunski, R. Patani, S. Chandran, G. Rot, B. Zupan, C. E. Shaw and J. Ule (2011). “Characterizing the RNA targets and position-dependent splicing regulation by TDP-43.” Nature neuroscience 14(4): 452–458.

Velasco, G., S. Grkovic and S. Ansieau (2006). “New insights into BS69 functions.” The Journal of biological chemistry 281(24): 16546–16550.

Wegorzewska, I., S. Bell, N.J. Cairns, T. M. Miller and R. H. Baloh (2009). “TDP-43 mutant transgenic mice develop features of ALS and frontotemporal lobar degeneration.” Proceedings of the National Academy of Sciences of the United States of America 106(44): 18809–18814.

Wen, H., Y. Li, Y. Xi, S. Jiang, S. Stratton, D. Peng, K. Tanaka, Y. Ren, Z. Xia, J. Wu, B. Li, M. C. Barton, W. Li, H. Li and X. Shi (2014). “ZMYND11 links histone H3.3K36me3 to transcription elongation and tumour suppression.” Nature 508(7495): 263–268.

Yu, B., Y. Shao, C. Zhang, Y. Chen, Q. Zhong, J. Zhang, H. Yang, W. Zhang and J. Wan (2009). “BS69 undergoes SUMO modification and plays an inhibitory role in muscle and neuronal differentiation.” Exp Cell Res 315(20): 3543–3553.

